# Stress induces distinct social behavior states encoded by the ventral hippocampus

**DOI:** 10.1101/2025.06.06.658117

**Authors:** Kevin Sattler, Michael Conoscenti, Addison Hedges, Jason Ortega, Raina Miller, Eva Owen, Ryan Hanson, Moriel Zelikowsky

**Affiliations:** Department of Neurobiology, University of Utah, School of Medicine. Salt Lake City, UT

**Keywords:** Stress, Aggression, Social hesitancy, Ventral Hippocampus, Trauma, PTSD

## Abstract

A single, acute traumatic experience can result in a host of negative impacts on behavior, such as increased violence, reduced sociability, and exaggerated fear responses. Despite the large body of research on the neurobiology of stress, we have a poor understanding of how trauma rewires social circuits in the brain. To identify how social circuits are re-organized by stress, we interrogated the role of the ventral hippocampus (VH), a key node for both the orchestration of emotionally-relevant behavior and the integration of sensory information. Using a footshock-based model of traumatic stress, we established that a single exposure to a series of unpredictable, inescapable footshocks was sufficient to negatively alter social behavior, resulting in increased violence and social hesitancy. Critically, we found that stress-induced changes to social behavior engaged neural ensembles in the VH, a critical node in the regulation of emotion states. Using a virally-mediated, activity-dependent cellular tagging approach to label two neural ensembles activated by temporally distinct experiences, we were surprised to find that stress-induced violence and stress-induced social hesitancy recruited largely non-overlapping populations of cells in the VH, in contrast to higher degrees of ensemble overlap in unstressed control mice, suggesting that stress biases the brain towards stronger differentiation of distinct social states. Additionally, we found that activity of VH neurons was required for stress-induced aggression. Finally, we found that stress-induced changes to social behaviors and VH activity profiles could be reversed by the introduction of social buffering post-stress. Collectively, our findings suggest that stress drives the VH to dissociably encode specific behavioral states, rather than a single state of negative valence, consistent with a role for multiple structures and distributed ensembles in the modulation of traumatic stress.

## INTRODUCTION

A single episode of traumatic stress has a profound ability to change our brain and behavior, resulting in a host of deleterious effects that can persist across the lifespan ^1,2^. Identification of the neurobiological mechanisms underlying the impact of stress to sensitize the brain and alter behavior has remained at the forefront of research efforts in recent years ^3–9^. These efforts have been fueled by rising levels of stress-related disorders, such as post-traumatic stress disorder (PTSD)^10,11^, the development of stress paradigms in model systems ^12–16^, and advances in circuit-based tools for the identification, manipulation, and recording of neural ensembles that drive stress-induced changes in behavior.

Footshock-based rodent models of stress provide a powerful tool to reveal the effects of stress. Exposure to footshock stress leads to wide-ranging and persistent changes to adaptive behavior, including increased fear sensitization ^17–19^, anxiety-like behavior ^20–25^, and depression-like behavior ^26–31^. However, relatively few studies have examined the impact of footshock stress to negatively alter subsequent social behavior, despite the increase in violence ^32^ and social withdrawal ^33^ that is known to occur in patients with PTSD. The handful of studies that have examined the impact of footshock stress on social behavior have utilized “double-hit” stress procedures, where footshock is combined with additional stressors, such as restraint stress, social defeat stress, and/or extended social isolation ^16,34–37^. While these models have helped to advance our understanding of the neurobiology underlying stress more broadly, they fail to elucidate the unique impact of a single stressful event on subsequent social behaviors. Thus, we adapted a model of footshock stress to identify the impact of a single, acute episode of footshock stress on subsequent social behaviors.

The ventral hippocampus (VH) is uniquely suited to modulate stress-altered social behavior, as it has been implicated in many different aspects of emotional processing, including fear and anxiety ^38–42^. More recently, findings have extended a role for the VH in social behavior ^43,44^, including mixed-stress-induced aggression ^37^, and deficits in social memory ^45,46^. This work suggests that the VH may be well suited to integrate changes in emotion state to drive alterations in social behavior. This idea of state integration is consistent with recent work demonstrating that anxiety-like behaviors induced by footshock stress are controlled by the VH ^47^.

Here, we performed a series of experiments aimed at tracking and perturbing neural ensembles in the ventral hippocampus (VH) during stress-altered social behavior. We found that acute footshock stress is sufficient to induce long-lasting and extinction resistant increases in fear, aggression, and social hesitancy in mice. Using a virally mediated, activity-dependent tagging system to identify changes to VH ensemble activity following stress, we found that distinct footshock-induced behaviors – aggression and social hesitancy – recruit dissociable neural ensembles in the VH and that chemogenetic inhibition of the VH attenuates stress-enhanced aggression. Positive social interactions can improve the deleterious effects associated with trauma in humans ^48–51^. Recent work has begun to highlight the positive impact of social buffering as a stress mediator in model organisms ^52,53^. Here we show that stress-induced changes to social behavior and VH activity were reversed by exposure to social conspecifics after stress, or “social buffering”. These findings support a role for the VH in the encoding of distinct social states induced by footshock stress and highlight a role for social buffering in the mitigation of social stress states.

## RESULTS

### Footshock stress produces enhanced fear learning in mice

To examine the impact of acute traumatic stress on subsequent social behavior in mice, we first adapted and validated a well-established model of footshock-based stress used in rats, stress-enhanced fear learning (SEFL) ^17^. While various models of traumatic stress have been developed for use in rodents, including mice, these often combine multiple stressors ^54^, making it difficult to determine whether subsequent changes in behavior are a result of a sensitized brain, or the additive effect of multiple stressors. We chose to adapt SEFL, as this rat-based model successfully recapitulates many of the behavioral phenotypes of PTSD ^17,18,21,29,55–61^. Recent versions of SEFL have demonstrated its potential ability to be translated for use in mice ^35,62–66^. Here, we develop a modified version of SEFL for use in mice, in which all additional stressors are omitted. In particular, we eliminate the confound of prolonged social isolation, as we have previously shown that prolonged isolation alone leads to wide-ranging and persistent impacts on social behavior ^67^.

Male mice received 10, pseudo-randomly delivered, unsignaled footshocks (FS; 1 sec, 1mA) or no footshocks (CTRL) over a 60-minute session in a novel context (Context A, Fig. 1A). The impact of this footshock stress to enhance subsequent fear learning was assessed by delivering a single mild footshock (2 sec, 0.35mA) in a novel context (Context B), and measuring subsequent contextual fear 24 hours later. We found that either a stronger mild shock or the delivery of more footshocks during stress produced ceiling effects in freezing for control mice (Fig. S1A-C), making it difficult to detect SEFL. To reduce baseline freezing after footshock stress, all mice were pre-exposed to Context B for 7 days prior to the mild footshock episode. Lack of context pre-exposure led to high levels of generalized freezing during the baseline period prior to mild shock (Fig. S1A-C).

**Figure 1.**
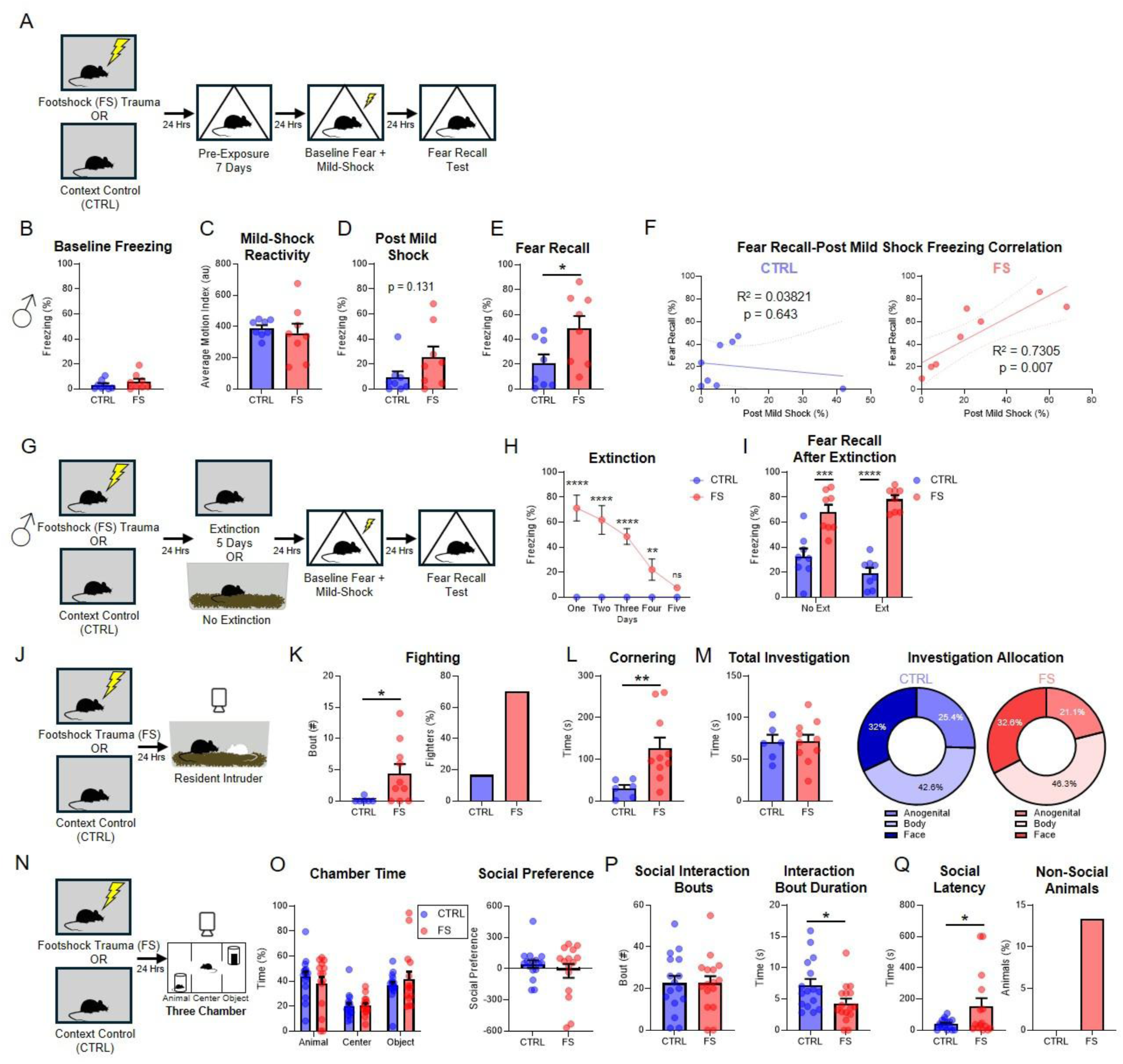
Footshock (FS) trauma produces enhanced fear learning in male mice and increases aggression and anxiety-like behaviors. (A) Experimental outline for stress-enhanced fear learning. (B) Male baseline freezing in Context B (C) Male reactivity to mild footshock during fear learning. (D) Male post-shock freezing. (E) Male freezing during fear recall test. (F) Male correlation between post-shock and fear recall test freezing in NS (left) and FS (right) mice. (G) Experimental outline for fear extinction SEFL experiment. (H) Percent of time spent freezing during extinction training. (I) Percent of time spent freezing during fear recall test. (J) Experimental design for resident intruder (RI) assay. (K) Fighting bouts during RI assay (left) and percentage of fighters (right). FS group also contains a higher percentage of animals that exhibit fighting behaviors. (L) Cornering behavior during RI assay. (M) Total time spent investigating intruder animal during RI assay (left) and percent allocation of anogenital, body, and face allocation for CTRL (middle) and FS (right). (N) Experimental design for the 3-chamber sociability assay (3Ch). (O) Percent time spent in each chamber during 3Ch assay (left) and social preference score (Animal Chamber Time - Object Chamber Time) (right). (P) Bouts of interaction with the novel conspecific (left) and duration of each individual social interaction (right). (Q) Latency to approach the novel conspecific (left) and percentage of animals that never approach the conspecific (right). Data are presented as mean ± SEM. Significant differences are reported as ****p < 0.0001, ***p < 0.001, **p < 0.01, and *p < 0.05.

Following footshock stress, male FS mice (n=8) exhibited similar levels of baseline freezing (Fig. 1B, unpaired t-test, t(14)=0.9909, p=0.339) and mild shock reactivity (Fig. 1C; unpaired t-test, t(14)=0.5071, p=0.620) when compared to CTRL mice (n=8). Additionally, FS mice showed no change in post-mild shock freezing (Fig. 1D; unpaired t-test, t(14)=1.605, p=0.131). We found that FS mice exhibited significantly elevated levels of freezing during context fear recall when compared to CTRL mice (Fig. 1E; unpaired t-test, t(14)=2.266, p=0.040), recapitulating the SEFL effect in our modified procedure. These results are consistent with the effects of stress to enhance subsequent fear learning and memory ^17,18,21,35,55,57,58,68–70^. To determine whether freezing immediately after the mild shock could be indicative of subsequent enhanced fear recall, we examined the correlation of percent time spent freezing between these two tests (Fig. 1F). We found that freezing levels post-mild shock and across fear recall were correlated in the FS, but not CTRL group (Pearson correlation, FS: r(6)=0.8547, p=0.007,CTRL: r(6)=-0.1955, p=0.643).

To determine whether our footshock stress protocol is sufficient to generate SEFL in females, female mice (CTRL, n=10; FS, n=10) were subjected to the same footshock stress protocol as males (Fig. S1D-H). Despite extensive context B pre-exposure, female FS mice showed increased baseline freezing compared to controls (Fig. S1D, unpaired t-test, t(18)=2.214, p=0.040), suggesting that females are more likely to exhibit persistent context fear generalization. Like males, FS females showed no change in shock reactivity (Fig. S1E, unpaired t-test, t(18)=1.640, p=0.118). They displayed increased freezing post-mild shock when compared to CTRL mice (Fig. S1F; unpaired t-test, t(17)=3.462, p=0.003). However, this enhancement was not maintained at recall test, as CTRL mice demonstrated high levels of context fear that were not different than FS mice (Fig. S1G, unpaired t-test, t(18)=0.5995, p=0.556). Interestingly, freezing levels during post-mild shock and fear recall tests showed no correlation in either CTRL or FS groups (Fig. S1H, Pearson correlation, CTRL: r(8)=0.601, p=0.066, FS: r(8)=0.5440, p=0.104). These data reveal that female mice expressing high levels of enhanced fear during the recall test showed variable levels of fear post mild-shock, suggesting that fear expression at test could not be predicted by how much a female froze post-shock. While other SEFL studies showed no significant sex differences using footshock combined with isolation in rats ^59,69^ or in a 129S6/SvEvTac mouse strain ^66^, our data suggest that inducing a fooshock-only stress state in C57Bl6/N female mice likely requires different experimental parameters than males.

### Enhanced fear is resistant to extinction

One of the hallmarks of trauma disorders is a resistance to extinction ^71^. This has been demonstrated in both humans ^72^ and rodents ^17,18,69,70^. To determine whether our footshock-based traumatic stress protocol is resistant to extinction, we introduced a series of extinction sessions after footshock stress and assessed subsequent enhanced fear (Fig. 1G-I). Male mice received either 5 days of extinction in Context A (Ext, n=8), or no extinction training (No Ext, n= 8; Fig. 1G). FS mice exhibited elevated levels of freezing during the first four days of extinction which were reduced by Day 5 (Fig. 1H, mixed-design ANOVA, Stress x Trial: F (4, 40) = 24.60, p<0.0001, Sidak post-hoc comparisons-Trials 1-4 p<.05, Trial 5 p = 0.650). After extinction, mice were subjected to mild footshock and context fear recall testing. We found that FS mice exhibited elevated freezing at recall compared to CTRL mice, regardless of their extinction condition (Fig. 1I, two-way ANOVA, Main Effect of Stress: F (1, 28) = 86.24, p<0.0001). These results suggest that the state induced by footshock stress is long lasting and extinction resistant in mice, consistent with previous work in rats ^17,18,20,69^.

### Footshock stress enhances fighting and social hesitancy

One of the hallmark symptoms of PTSD is an increase in anger ^32^, social anxiety ^33^, enhanced startle responses ^73,74^, and depression ^75^. Nonetheless, the effect of traumatic stress to negatively alter social behavior has remained largely understudied in model systems. To examine the impact of footshock stress on social behavior in mice, we subjected mice to our footshock stress protocol and tested them for subsequent changes to social behavior using the Resident Intruder (RI) assay to assess aggression ^67,76^ and the 3-Chamber social preference assay to assess social interactions ^77^.

Male mice were first tested on the RI assay following exposure to footshock stress (FS), or spent equivalent time in the footshock context with no shock exposure (CTRL) (Fig. 1J). Behavior captured during the RI assay revealed increased fighting in FS mice (n=10) compared to CTRL mice (n=6; Mann-Whitney U=10.50, p=0.0237). Additionally, a higher percentage of animals in the FS group engaged in fighting behavior compared to controls (Fig. 1K). In a separate cohort of mice, we found that this stress-induced aggression was resistant to extinction (Fig. S1I,J), demonstrating that stress-induced aggression, like stress-enhanced fear, is indicative of a PTSD-like state. In addition, RI testing revealed an increase in social cornering behavior for FS mice (Fig. 1L, unpaired t-test, t(14)=2.806, p=0.014). We observed no changes in total social investigation time (unpaired t-test, t(14)=0.06745, p=0.947) or in how they allocate the amount of time spent investigating each subregion of the conspecific intruder mouse (unpaired t-tests, Anogenital: t(14)=0.7937, p= 0.441, Body: t(14)=0.8106, p=0.431, Face: t(14)=0.1172, p=0.908; Fig. 1M). These data demonstrate that footshock stress generates a sensitized state in which social behaviors, such as aggression, are enhanced.

While the RI assay allows for unrestricted social interaction, it does not probe social motivation, which has been shown to be negatively impacted in patients with PTSD ^78^. To test whether our model of footshock stress impacts subsequent social preference and motivation in mice, we used the 3-chamber sociability assay (3Ch) ^77^; Fig. 1N). FS and CTRL mice were tested for sociability in the 3Ch for 10 minutes, where a novel sex-, age- and strain-matched conspecific mouse was placed in one chamber (“social”), a novel object was placed on the opposite chamber (“object”), and the center chamber remained free of any stimuli. We found that both FS and CTRL mice spent more time in the social and object chambers than the middle chamber (two-way ANOVA, Main Effect of Chamber, F (2, 87) = 16.76, p<0.001, Tukey’s posthoc analysis, Social vs. Object: p= 0.911, Social vs. Middle: p<0.001, Object vs. Middle: p<0.001). Interestingly, we found no social preference (time spent in social chamber-time - spent with object) for either CTRL (n=16) or FS (n=15) mice (unpaired t-test, t(29)=0.8505, p=0.402; Fig. 1O). However, this finding is consistent with a general variability in whether stress impacts social preference in the 3 Chamber assay ^79–81^. However, while there was no change in the number of social interaction bouts between groups (unpaired t-test, t(30)=0.000, p>.999), the average duration of each bout was significantly shorter in FS mice (unpaired t-test, t(30)=2.295, p=0.029; Fig. 1*P*, Video S1 and S2). Moreover, the latency for FS mice to first approach conspecific mice was significantly longer (unpaired t-test, t(29) = 2.107, p=0.0439), and a subset of FS mice never socialized with the conspecific at all (Fig. 1Q), revealing increased social hesitancy in FS mice. These data demonstrate that footshock stress changes the nature of social interactions by increasing social hesitancy and reducing the time spent interacting socially. As with SEFL, behavior of FS female mice on both the RI and 3Ch assays was not different then controls (Fig. S1O-W), again suggesting that females likely require distinct, sex-specific parameters to generate a state of footshock stress.

### Footshock stress-induced aggression engages the VH

Our data reveal that footshock stress induces a sensitized state, which can lead to deleterious impacts on fear and social behavior, including enhancing aggression and social hesitancy. Recent findings have shown that the sensitized state induced by stress is encoded across numerous limbic structures, including the ventral hippocampus (VH) ^21,65,82,83^. In a separate set of studies, the VH was found to modulate increased aggression following social isolation via projections to the hypothalamus ^84^. These findings, combined with the recent identification of the VH in the control of footshock stress-induced anxiety ^47^, suggest that the VH may play a privileged role in encoding stress states and motivating subsequent changes in behavior.

To determine whether the VH is activated following FS-altered social behavior, we first performed an activity-dependent labeling experiment. Mice were subjected to our FS (n=3) or CTRL (n=3) protocol and tested for social behavior using the RI assay. Brains were extracted and processed for cFos expression in the ventral CA1 (vCA1) subregion of the VH, as the vCA1 has been shown to project directly to the ventromedial hypothalamus to modulate isolation-induced aggression ^84^. We found that cFos expression in the VH was increased in FS compared to CTRL mice after the RI assay (unpaired t-test, t(3) = 4.818, p=0.0170; Fig. 2A, B), suggesting that the VH is activated by stress-induced changes to social behavior during free interaction with an intruder mouse. Importantly, we did not see FS-increased cFos expression in the dorsal CA1 (t(5)=0.1264, p=0.9044; Fig. 2C,D), consistent with literature suggesting that the ventral subregion of the VH is preferentially tuned to stress ^85,86^.

**Figure 2.**
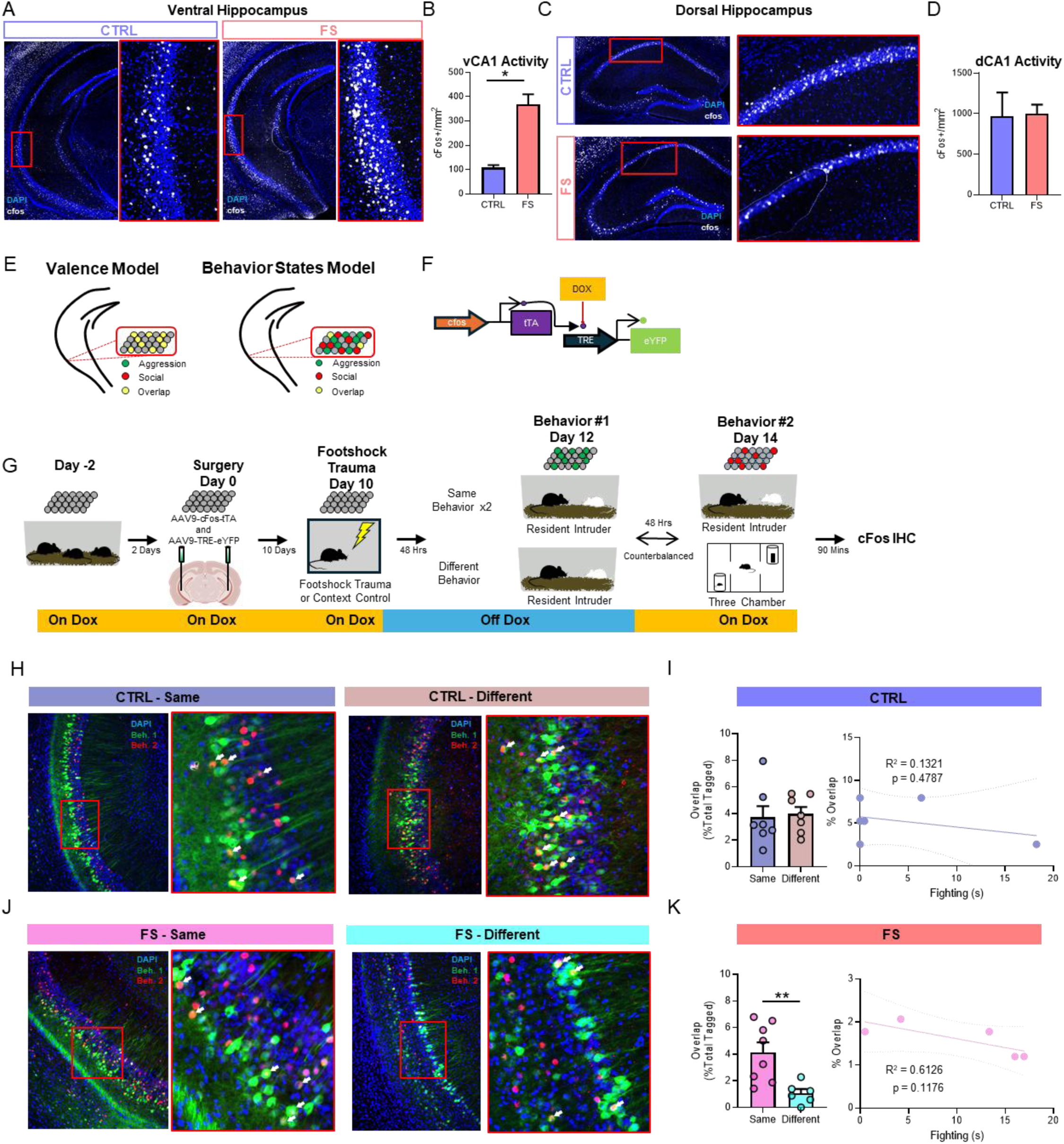
Dissociable stress-induced behavior states are encoded by distinct ensembles in the ventral hippocampus. (A) Representative images for VH cFos upregulation in CTRL (left) and FS (right). (B) Quantification of cFos activity in the vCA1. ***C.*** Representative images for DH cFos activity in CTRL (top) and FS (bottom). (C) Representative images for DH cFos activity in CTRL (top) and FS (bottom). (D) Quantification of cFos activity in the dCA1. (E) Proposed models of VH modulation of trauma altered social behaviors. Valence model (left) would suggest that all cells are trauma cells and modulate before equally. Behavior states model (right) would suggest that VH neurons are behavior specific, not valence specific. (F) Schematic representation of the activity-dependent tagging viruses. (G) Experimental design to tag active neurons in the VH during different behavioral protocols. (H). Representative images for CTRL animals after experiencing the same behavioral protocol twice (left) or different behavioral protocols (right). (I) Quantification of cellular overlap (left) and correlation graph showing no correlation between fighting and cellular overlap (right) in control mice. (J). Representative images for FS animals after experiencing the same behavioral protocol twice (left) or different behavioral protocols (right). (K) Quantification of cellular overlap (left) and correlation graph showing no correlation between fighting and cellular overlap (right) in FS mice. (L) Cornering behavior during RI assay. (M) Total time spent investigating intruder animal during RI assay (left) and percent allocation of anogenital, body, and face allocation for CTRL (middle) and FS (right). (N) Experimental design for the 3-chamber sociability assay (3Ch). (O) Percent time spent in each chamber during 3Ch assay (left) and social preference score (Animal Chamber Time - Object Chamber Time) (right). (P) Bouts of interaction with the novel conspecific (left) and duration of each individual social interaction (right). (Q) Latency to approach the novel conspecific (left) and percentage of animals that never approach the conspecific (right). Data are presented as mean ± SEM. Significant differences are reported as ****p < 0.0001, ***p < 0.001, **p < 0.01, and *p < 0.05.

### Stress induces the encoding of distinct behavior states by separable ensembles in the VH

Our data demonstrate that footshock stress negatively alters social behavior, and that these effects correlate with increased activity in the VH. However, little is known about whether such sensitized states control behavioral changes via a single neural ensemble, which represents a negatively-valenced state, or whether dissociable ensembles in a given region are recruited to drive distinct behaviors produced by that state. We tested whether ensemble activity in the VH tracks the general negative valence of a stress state (“valence model”), or the distinct behavior states being induced by stress (“behavior states model”) (Fig. 2E).

To determine whether footshock stress-induced aggression and social hesitancy are encoded by a single, unified population of cells in the VH, or by distinct subpopulations of cells, we took an activity-dependent dual labeling approach using a viral cocktail-mediated, activity-dependent strategy for cell tagging ^87^. The viral cocktail strategy we utilized (Fig. 2F) – AAV9-cFos-tTA + AAV9-TRE-eYFP – allows for the cFos promoter to drive expression of tTA (tetracycline-controlled transactivator), which will then bind to TRE (tetracycline response element) and drive the expression of a fluorophore for fluorescent tagging (i.e., eYFP). This system is under the control of doxycycline (DOX), which, when present, prevents the binding of tTA to TRE. We combined this approach with immunohistochemistry for cFos to allow for visualization of two activity-dependent cellular tagging epochs within the same mouse.

Mice were placed on DOX and injected with the cFos-tTA + TRE-eYFP viral cocktail into the VH (Fig. 2F,G) before being subjected to FS-stress or context control exposure. Immediately following FS or context exposure, mice were taken off DOX to open the first tagging window. Mice were then subjected to either the RI or 3Ch assay (first tag) and immediately placed back on DOX to close the tagging window for assay 1. Forty-eight hours later, mice were subjected to a second behavioral assay (either the same behavioral assay (RI→RI or 3Ch→3Ch) or the opposite assay (RI→3Ch or 3Ch→RI), after which brains were extracted and processed for cFos (second tag) (Fig. 2G). Therefore, cells activated by the first behavioral assay were tagged with eYFP (green) and cells activated by the second behavioral assay were tagged immunohistochemically (red). These conditions allowed us to determine whether cFos expression profiles showed significant overlap across two distinct stress-induced behaviors, aggression and social hesitancy, as compared to the degree of overlap seen following exposure to the same assay twice.

CTRL mice showed no difference in the percentage of behavior-labelled cells with overlap in the VH when animals experienced the same behavior twice (n=7) compared to different behaviors (n=7; Fig. 2H, I); unpaired t-test, t(12)=0.2480, p=0.808), consistent with the general idea that the DH, not the VH, encodes contextual information. In addition, the degree of ensemble overlap was not correlated with the amount of fighting behavior observed (Fig. 2I,K), further highlighting that under normal conditions, an animal’s behavior in a given environment does not drive differential encoding by the VH. Surprisingly, FS mice showed a significant decrease in the percentage of ensemble overlap following different behaviors (n=6) compared to the same behavior twice (n=8; Fig. 2J, K; unpaired t-test, t(12)=3.281, p=0.007), suggesting that in a stressed state, the VH becomes more highly tuned to environmental and behavior conditions, and that distinct behaviors become encoded by dedicated, largely non-overlapping populations of neurons in the VH. Collectively, these data suggest that under normal conditions, the VH may broadly encode social behavior without behavioral specificity but that under states of stress, the VH recruits dissociable ensembles of neurons for distinct behaviors. These findings suggest that footshock stress promotes the encoding of distinct behavior states, not valence, in VH ensembles.

### Chemogenetic silencing of the VH attenuates footshock-induced aggression

To determine if the VH not only correlates with but is necessary for FS-induced changes in social behavior, we used a chemogenetic approach to inhibit the VH during the RI (Fig. 3A) or the 3-Chamber assay (Fig. 3E) following FS. Mice were injected with a virus (AAV2-hSyn-hM4Di) encoding the inhibitory DREADD, hM4Di ^88^, into the VH. Following recovery, all mice underwent footshock stress followed by testing on the RI (n=29) or 3Ch assay (n=10). Mice were tested on the same assay twice – once following injection of the DREADDs-activating ligand DCZ (deschloroclozapine, 1mg/kg, ip) ^89,90^ and once following a saline vehicle injection (VEH; order counterbalanced), to enable within-subjects comparisons.

**Figure 3.**
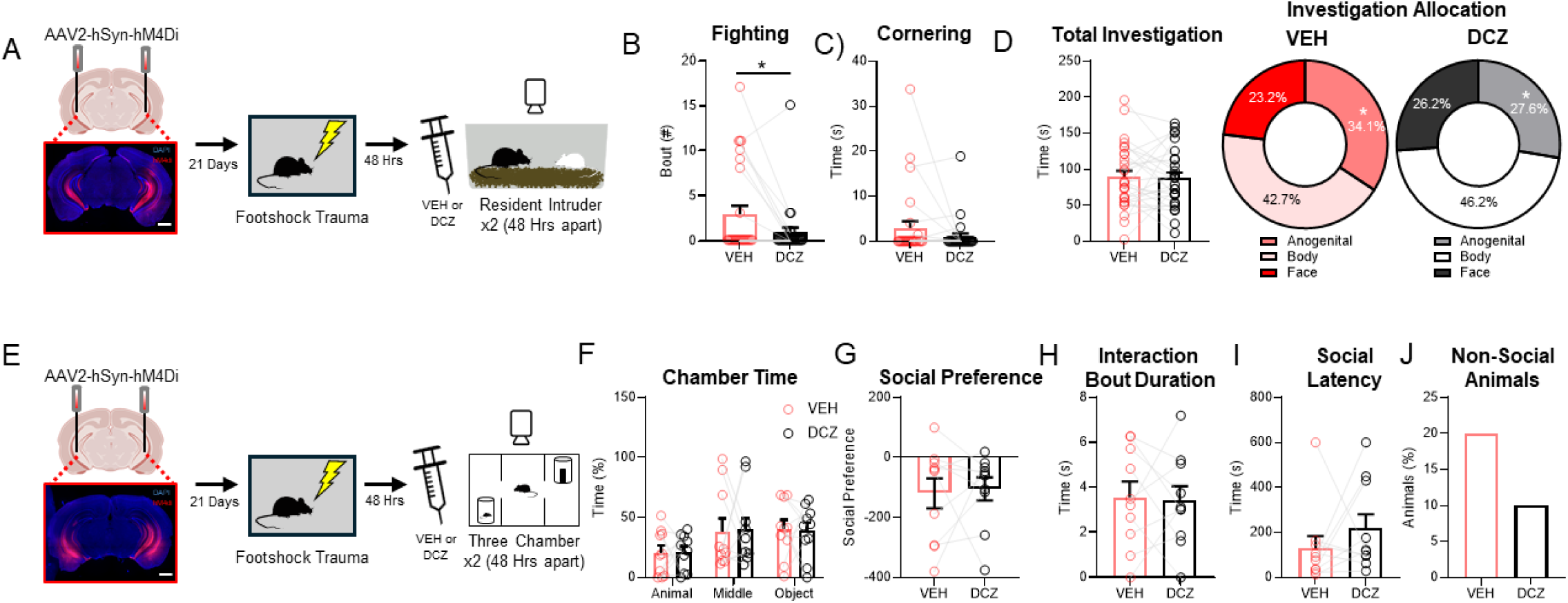
Chemogenetic silencing of the VH ameliorates footshock enhanced aggression. (A) Experimental design and representative image for chemogenetically silencing the VH during RI assay. (B) Fighting bouts during RI with the VH active (VEH) and the VH silenced (DCZ). (C) Cornering time during RI with the VH active (VEH) and the VH silenced (DCZ). (D) Total time spent investigating intruder animal during RI assay (left) and percent allocation of anogenital, body, and face allocation for CTRL (middle) and FS (right) with the VH active (VEH) and the VH silenced (DCZ). (E) Experimental design and representative image for chemogenetically silencing the VH during the 3-chamber sociability assay. (F) Percent time spent in each chamber during 3Ch assay (left) and social preference score (Animal Chamber Time - Object Chamber Time) (right) with the VH active (VEH) and the VH silenced (DCZ). (G) Duration of each individual social interaction with the VH active (VEH) and the VH silenced (DCZ). (H) Latency to approach the novel conspecific (left) and percentage of animals that never approach the conspecific (right) with the VH active (VEH) and the VH silenced (DCZ). Data are presented as mean ± SEM. Significant differences are reported as **p < 0.01, and *p < 0.05.

We found that silencing the VH significantly decreased fighting bouts in DCZ-injected mice (paired t-test, t(27)=2.491, p=0.019; Fig. 3B). In contrast, silencing the VH had no effect on cornering behavior (paired t-test, t(27)=1.406, p=0.171;Fig. 3C) or total investigation time (paired t-test, t(27)=0.2241, p=0.824;Fig. 3D, left). While there was no change in total investigation time, silencing the VH decreased the percentage of time mice spent investigating the anogenital region of the intruder mouse (paired t-test, t(28)=2.3274, p=0.0274; Fig. 3D, right, Fig. S3A-C). These results suggest that the VH is required for FS-induced aggression and may play a further role in mediating the impact of FS to induce more subtle changes in social behavior.

In contrast, when we inhibited the VH during the 3Ch sociability assay, there was no effect on time spent in each chamber, social preference, social interaction bout duration, or social latency (Fig. 3F-I; chamber time: unpaired t-tests, animal chamber: t(18)= 0.0865, p=0.9320, middle chamber: t(18)=0.0829, p=0.9348, object chamber: t(18)=0.1842, p=0.8559; social preference: unpaired t-test, t(18)=0.2388, p=0.8139; social interaction: unpaired t-test, t(18)=0.1585, p=0.8758; social latency: unpaired t-test, t(18)=1.068, p=0.2995), although the number of mice that never entered the social chamber was reduced (Fig. 3J). These data reveal that although the VH contains neural ensembles that are activated by distinct footshock-induced behaviors, activity in this region may only be required for some but not all the effects induced by FS.

### Social buffering ameliorates trauma-enhanced aggression

Social support or “buffering” is emerging as a potent mitigator of trauma-induced negative changes to behavior ^48–53,91–93^. To determine whether social exposure can influence footshock-induced changes in social behavior, we updated our behavioral protocol to include a period of social conspecific exposure (“social-buffering”) following stress (Fig. 4A). As in our previous experiments, mice were either subjected to footshock stress or control context exposure (CTRL). FS mice were then immediately split up and placed into 1 of 3 housing conditions: housed alone (“no social buffer” group; FS+NB), co-housed with mice whom had also received footshock (“collective trauma” group; FS+CT), or co-housed with naïve, non-stressed mice (“social buffer” group; FS+SB) (Fig. 4A). Following 48 hours of post-stress housing, mice were tested on the RI assay. Consistent with our original findings (Fig. 1), footshock stress (FS+NB, n=20) increased the number of fighting bouts when compared to control mice (CTRL, n=11). This effect was decreased by co-housing with stressed mice (FS+CT, n=14) and eliminated when the mice were co-housed with naïve, non-stressed mice (FS+SB, n=13) (Fig. 4B *left*, One-way ANOVA: f(3,54)=3.634, p=0.0184; Holm-Sidak’s multiple comparisons: FS+NB vs CTRL, p=0.0453; FS+NB vs FS+CT, p=0.0716; FS+NB vs FS+SB, p=0.0288). The percentage of animals that engaged in attack was also abolished in mice co-housed after stress (Fig. 4B *right*). The same pattern of effects was observed for cornering behavior (Fig. 4*C*, One-Way ANOVA: f(3,54)=4.567, p=0.0064; Holm-Sidak’s multiple comparisons: FS+NB vs CTRL, p=0.0261; FS+NB vs FS+CT, p=0.0762; FS+NB vs FS+SB, p=0.0079). These data suggest that social contact following stress, particularly with naïve conspecifics, serves as a social buffer to mitigate against the effects of stress to induce aggression and cornering behavior.

**Figure 4.**
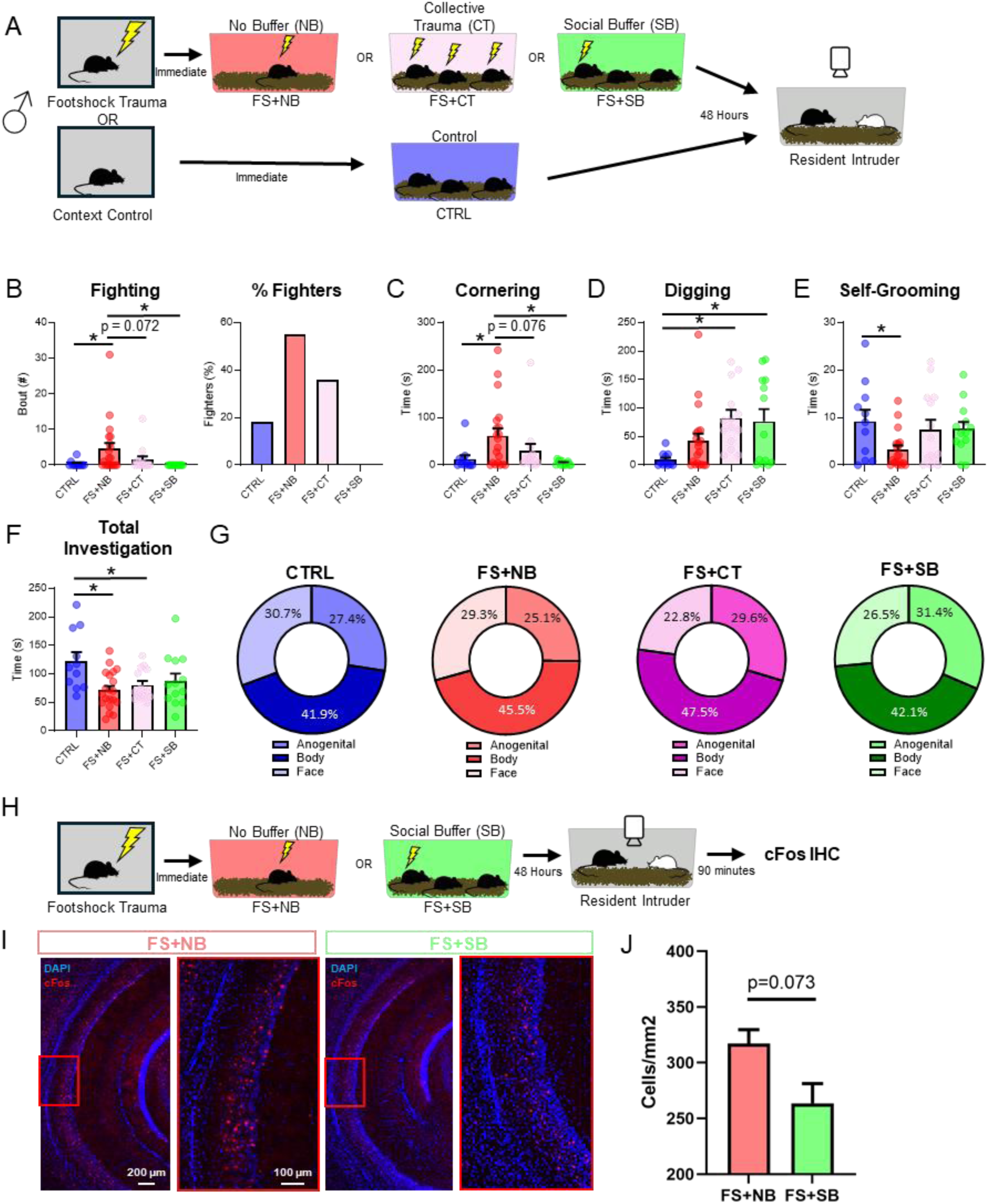
Social buffering is sufficient to ameliorate trauma-enhanced aggression and improve self-care and anxiety-like behaviors but has no effect on deficits in prosocial investigation. (A) Experimental design for social buffering and resident-intruder. (B) Fighting bouts during RI assay (left) and percentage of fighters (right). FS group also contains a higher percentage of animals that exhibit fighting behaviors. (C) Cornering behavior during RI assay.. (D) Digging behavior during RI assay. (E) Self-Grooming behavior during RI assay. (F) Total time spent investigating intruder animal during RI assay (combined anogenital, body, and face investigation). (G) Percent allocation of anogenital, body, and face investigation for each experimental group. Data are presented as mean ± SEM. Significant differences are reported as **p < 0.01, and *p < 0.05. (H) Experimental design for VH cFos quantification. (I) Representative images for VH cFos quantification in FS+NB (left) and FS+SB (right). (J) Quantification of cFos activity in the vCA1.

We also examined non-social behaviors during the RI assay, including digging – a behavior associated with elevated anxiety ^94^, and self-grooming – a behavior associated with self-care ^95^. Interestingly, we found a significant increase in the amount of digging produced in both social co-housing groups (Fig. 4*D*, One-Way ANOVA: f(3,54)=5.542, p=0.0022; Holm-Sidak’s multiple comparisons: CTRL vs. FS+CT, p=0.0124; CTRL vs FS+SB, p=0.0245), suggesting that while social buffering is beneficial for certain forms of anxiety-like behavior (e.g. cornering), it may exacerbate others, or, that digging is a complex behavior that remains to be thoroughly explored. Alternatively, social buffering may shift the mice from an aggressive state to a state of general arousal, thereby decreasing attack behavior while simultaneously increasing behavior such as digging. Interestingly, we found that self-grooming- a behavior in mice involved in hygiene and thermoregulation ^95^- was significantly decreased following footshock stress, but not for animals that had undergone co-housing of any kind (Fig. 4E, One-way ANOVA: f(3,53)=2.992, p=0.0390; Holm-Sidak’s multiple comparisons: CTRL vs. FS+NB, p=0.0408; CTRL vs FS+CT, p=0.7444; CTRL vs FS+SB, p=0.7444), suggesting that social buffering can partially recover deficits in self-care following footshock exposure. Lastly, footshock caused a significant decrease in overall time spent on investigation behaviors, an effect that was not reversed by co-housing (Fig. 4F, One-way ANOVA: f(3,54)=1.314, p=0.0092; Holm-Sidak’s multiple comparisons: CTRL vs FS+NB, p=0.0058; CTRL vs FS+CT, p=0.0492; CTRL vs FS+SB, p=0.1347). There were no significant differences in how mice allocate the time spent investigating the different regions on the intruder mouse (Fig. 4*G*, One-way ANOVAs, anogenital: F(3,54)= 1.012, p=0.3945, body: F(3,54)= 1.025, p=0.3890, face: F(3,54)= 1.081, p=0.3649).

To determine whether the behavioral effect of social buffering correlate with changes in VH activity, we tested whether social buffering could reverse footshock-induced increases in VH activity. All mice were exposed to footshock and received either no social buffering (FS+NB, n=4) or social buffering (FS+SB, n=2) and were assayed in the RI box 48-hours later (Fig. 4H). Ninety minutes following the RI assay, mice were perfused and brains were extracted for IHC quantification of cFos. To control for confounds of aggression-induced VH activity, only non-aggressive mice were included in the analysis. We found that mice exposed to footshock and co-housed with unshocked companions exhibited a decrease in cFos expression compared to their non-socially buffered counterparts (Fig. 4I,J; nested t-test, t(4)=2.415, p=0.0732). These data suggest that the effect of social buffering to reverse many of the effects of footshock stress on behavior may be mediated by the ventral hippocampus.

## DISCUSSION

Traumatic stress negatively impacts the social fabric of our lives – disrupting inter-personal relationships, increasing domestic violence, promoting social anxiety and more ^33,96–98^. Despite the ability of acute trauma to dramatically alter social behavior, the neural mechanisms underlying this effect have been largely unexplored. Here, we show that exposure to footshock stress in mice leads to increased aggression and the promotion of social hesitancy. We identify a role for dissociable ventral hippocampal ensembles in the encoding of distinct social behaviors impacted by stress, consistent with the idea that the VH encodes distinct behavior states. We find that VH activity not only tracks the behavior state of mice but is necessary for footshock-induced aggression. Lastly, we show that post-stress social buffering significantly ameliorates the behavioral effects of stress and decreases VH neuronal activity.

### Capturing sex differences in models of PTSD

One critical result we obtained is that our adaptation of a well-validated model of PTSD in rodents did not successfully translate to females. This contrasts with prior research on SEFL in rodents, where similar effects in both male and females was found, though this was restricted to rats or non-traditional mouse strains ^59,62,66,69^. Clinically, women develop PTSD at a much higher rate than men ^99^ and report more severe and longer lasting pain with similar disease processes to men ^100^. Collectively, this literature suggests that if anything, footshock stress should produce even stronger PTSD-like effects in females than in males. However, studies in mice suggest that females exhibit greater variability in fear behaviors, including less sensitivity to footshock ^101^, but greater fear-potentiated startle ^102^. Thus, a potential explanation for the absence of enhanced fear learning and/or aggression in female mice in our study is that the footshock intensity (mA) used for the mild-shock was too strong, leading to high levels of freezing in control mice at test. This, combined with a ceiling effect in FS mice ^103^, could have prohibited us from identifying any potential SEFL effect in FS mice. This would be surprising, as female rodents tend to exhibit weaker fear acquisition when compared to their male counterparts ^20,104,105^. However, given that our study had significant context pre-exposure, the saliency and/or magnitude of the shock could explain this discrepancy. This suggests that sex interacts with the severity of stress to produce sex-specific outcomes, and that additional refining of our footshock stress parameters will be required to tailor our model for use in female mice.

### Disentangling the parameters of stress-induced aggression and sociability

Our results indicate that a single episode of footshock stress is sufficient to induce aggression in mice. These data are consistent with numerous findings which demonstrate an increase in aggression following stress ^16,35,37,106^. However, our results indicate that footshock stress alone is capable of inducing aggression, without the inclusion of additional stressors, such as prolonged social isolation, which is sufficient to cause aggression on its own ^80,107–109^. This is critical because it demonstrates that a single exposure to traumatic stress is sufficient to generate increased violence in male mice, a phenotype exhibited by a subset of patients with PTSD ^32^. Importantly, these data, in combination with numerous findings by others in the field of aggression ^110–118^, reveal that there are numerous environmental experiences which are sufficient to engender aggression. Future research is warranted to identify shared and unique mechanisms governing aggression induced by distinct conditions.

In addition to changes to social behavior in the RI assay, we found that stressed mice also demonstrated differences in social behavior during the 3 Chamber assay. Indeed, FS mice took longer to approach the social cup and had shorter bouts of social interaction, when they did make contact with the contained conspecific mouse. Strikingly, a small subset of FS mice never entered the social chamber at all. While we observed this social hesitancy, we did not see overall reductions in social preference for FS mice. These findings are consistent with known variability in social preference in the 3 Chamber following other forms of stress. For example, exposure to social isolation produces opposing effects on total time in the social chamber, depending on the length of the isolation period – more interaction after acute isolation, less interaction after chronic isolation ^67,119,120^. Meanwhile, chronic non-social stressors, such as chronic forced swim and chronic restraint stress, appear to consistently decrease social preference ^121–123^. In addition, we and others have found that the restrictive social nature of the 3 Chamber test may introduce limitations in the ability to examine real changes to sociability ^77,79,124–127^. Future studies using more naturalistic tests of sociability are warranted.

### The ventral hippocampus drives distinct behavior-state encoding following stress

The ventral hippocampus has repeatedly been identified as a key node in the neural circuitry controlling stress ^47^, anxiety ^41,128^, and negative valence ^39^. Most recently, a pair of findings has revealed a role for the VH in controlling anxiety-like behaviors after footshock stress ^47^ as well as anxiety ^129^. In addition, recent work has begun to expand our understanding of the VH as a modulator of social behaviors, including aggression ^37^ and social memory ^46^. In line with this literature, we found that stress-induced aggression engages the ventral hippocampus. We hypothesized that similar neural ensembles in the VH would be activated by various behaviors impacted by stress, consistent with the idea that the VH encodes negative valence states ^42^, such as that produced by footshock stress. Surprisingly, however, we found that neural ensembles in the VH that were activated by either FS-induced aggression or FS-induced social hesitancy were largely non-overlapping. This directly contrasts with our hypothesis that a unified ensemble representing the negatively-valanced state of footshock stress would be similarly active during each FS-induced social behavior. Instead, our data demonstrate that dissociable, non-overlapping ensembles in the VH encode each distinct FS-induced behavior states, suggesting a “behavior states” model for VH encoding rather than a “valence”-based model (Fig. 3E). While the percentages of overlap are low in both conditions, this is consistent with other VH literature using similar techniques ^130^. These findings challenge the idea of the VH as a valence detector ^40,131^ but are in line with recent research suggesting that the VH may exhibit some contextual or experiential functions similar to its dorsal counterpart ^132^ and that the VH may function more like the dorsal hippocampus under certain conditions ^133^. These findings also dovetail with recent literature demonstrating that models of PTSD-like behavior in rodents, such as SEFL, are encoded across distributed brain regions, rather than being controlled by a single hub region (Pennington et al., 2024).

### Social buffering mitigates against stress-altered social behavior

Social interaction has been shown to improve non-social PTSD-like symptoms in humans ^49^ and rodents ^53^. Here, we extend these findings to changes in social behavior. We show that social buffering via co-housing with conspecific mice, particularly those whom have not undergone stress themselves, is sufficient to reverse the effects of footshock on numerous behaviors, including the induction of aggression. These findings demonstrate the powerful impact that social contact and connection can have on the well-being of an animal. As trauma-related mental health disorders have been increasing over the past several years ^134^, these results are timely, suggesting potential points of therapeutic, non-invasive intervention following trauma.

One of the most debilitating impacts of PTSD is its ability to generate violence and destroy social relationships. Nevertheless, our understanding of the negative impact of trauma on social behavior has remained relatively sidelined in contrast to our understanding of how trauma enhances fear, depression, and anxiety. Our results present a behavioral model in which footshock-induced aggression and social hesitancy can be investigated, and the underlying neurobiology interrogated. We expect that these findings will move the field forward in our understanding of traumatic stress and the distinct social behavior states induced by stress.

## METHOD

### Animals

Wild-type males and females aged 10-12 weeks (Charles River Laboratories) were housed in groups of 4 per cage. All mice were kept on 12:12 light/dark cycles (10:00-22:00) in a humidity-controlled mice vivarium with *ad libitum* access to standard rodent chow and water. Upon arrival at the facility, mice were left undisturbed for one week to acclimate and recover from transport. For activity-dependent tagging experiments, we substituted the rodent chow with 40 mg/kg doxycycline (DOX) chow 24-hours prior to surgery and these animals were given 10 days to recover. For chemogenetic inhibition of the ventral hippocampus, mice were given 21 days following viral injection surgery to recover and allow time for the DREADDs to express. All subjects were treated in accordance with our protocol approved by the University of Utah’s Institutional Animal Care and Use Committee (IACUC).

### Activity Dependent Tagging

The two viruses for the activity-dependent tagging system were packaged at the University of Massachusetts Amherst Viral Vector Core and provided courtesy of Steve Ramirez. The first virus contains a pAAV-cFos-tTA plasmid vector and the second a pAAV-TRE-eYFP. Neural activity induces the expression of the AAV9-cFos-tTA construct and generates tTA proteins in cells. The tetracycline transactivator (tTA) proteins then bind to the tetracycline response element (TRE) to induce the expression of -eYFP. This system is regulated by DOX, a tetracycline derivative, for strict temporal control over when the cells are labeled. The removal of DOX from the system opens a tagging window to allow tTA to bind to TRE.

### Surgical Procedures

Mice were anesthetized in an induction chamber using 4-5% isoflurane and anesthesia was maintained at 1-2% isoflurane while the mice were mounted on the nose cone of the stereotaxic surgery rig (Kopf Instruments, California, USA). Ophthalmic ointment was applied to both eyes following stereotax mounting to prevent eyes from drying during surgery. A topical hair removal cream was used to remove hair on the scalp and the surgical area was cleaned 3x with isopropyl alcohol and betadine. Subcutaneous injection of buprenorphine (3.25 mg/kg) was given prior to any incisions as an analgesic. A small incision was made to expose the skull, the skull was leveled between bregma and lambda, and bilateral holes were drilled above the site of viral injections.

Injection coordinates into the ventral hippocampus are in relation to bregma (in mm). For DREADDs experiments, bilateral injections were performed at AP: −3.0, ML: +/-3.2, DV: −4.2. In tagging experiments, two injections were performed/side, in an attempt to cover the entirety of the ventral hippocampus (AP: −3.0, ML: +/-3.2, DV: −4.2 & −3.0). Virus was backfilled into pulled fine glass capillaries (∼ 50 um diameter at tip) and pressure injections of 100-250 nl were made bilaterally (250 nl for DREADDS, 200 & 100 nl for tagging) at a rate of 75 nl/min using a nanoliter injector (Nanoliter 2000, World Precisions Instruments) controlled by an ultra microsyringe pump (Micro4, World Precision Instruments). Capillaries remained in place for 5 minutes following injections to allow for full diffusion of virus and to reduce backflow up the injection tract. Skin above the skull was then drawn together and sealed with GLUture (Zoetis). All injections were subsequently verified histologically.

### Behavioral Assays

#### Footshock Stress and Fear Conditioning

Stress pretreatment and testing occurred in Med Associates behavioral testing chambers. Each chamber was equipped with an infrared camera, speaker for tone delivery, shock scrambler, and fluorescent and infrared light sources. The behavioral testing chambers in each testing room were controlled by a PC using Med Associates Video Freeze software that also automatically scores motion and freezing of the animal during the test session. A mouse is considered freezing when an image change is registered at less than 30 pixels for at least 1 second. Any baseline pixel change due to mechanical operation of the chamber or camera is measured before the animal’s entry and subtracted from measurement during the trial. To create distinct contexts between stress pretreatment and subsequent fear conditioning and testing, the chamber’s contextual features were modified using differential lighting, odors, ambient noise, grid floors, and wall inserts. Context A was performed in chambers during the light cycle with white lights on in the room and chamber, a floor with a single layer, fans in the chambers on, and the chambers were cleaned with 70% ethanol and scented with 50% Simple Green. Although performed at a similar time of day, Context B was performed with only red lights on in the room, infrared light only in the chamber, no fan noises, the floor was double layered, and the chambers were cleaned with 70% isopropyl alcohol and scented with Windex. Unless otherwise stated, contexts differed on as many dimensions as possible to reduce generalization effects.

All mice were moved into individual cages two days prior to stress pretreatment. Stress pretreatment included 10, 1 second, 1mA un-signaled footshocks pseudo-randomly distributed over a 60-minute session in a novel context (Context A). Control mice were exposed to the same context for sixty minutes but received no footshock. In experiments where subsequent fear learning was assessed, all mice were pre-exposed to a novel context (Context B) for 7 days prior to the mild footshock episode. During these sessions, mice were placed into the context for 30 minutes and allowed to explore freely. Following context pre-exposure, all mice received a single mild footshock (2-sec, 0.35mA) in Context B. Three minutes of context exposure preceded the shock, and mice were removed from the chamber 1 minute after the termination of shock. Baseline freezing is an average of freezing for the 3 minutes preceding the footshock. Shock reactivity was measured by recording the average motion index during the mild-shock and for the 3 seconds following the shock. The following day, mice were placed back into Context B for an 8-minute session, to test for contextual fear recall.

#### Resident Intruder Assay

Testing for aggression using the resident intruder assay (Blanchard et al., 2003) proceeded as previously described (Hong et al., 2014, 2015; Lee et al., 2014). Briefly, experimental mice (“residents”) were transported via cart from the vivarium to the behavioral testing suite and allowed to acclimate for at least one hour. At the time of testing, mice were transported in their homecage to a novel behavior testing room, from an adjacent “holding” room. Homecages were then slotted into a customized behavioral chamber lit with infrared lights and equipped with two synchronized infrared video cameras placed at 90-degree angles from each other to allow for simultaneous behavior recording with a front and top view. Synchronized video was acquired using Media Recorder software (Noldus, 30ms frame rate). Following a two-minute baseline period, an unfamiliar male BALB/c mouse (“intruder”) was placed in the homecage of the resident for 10 minutes and mice were allowed to freely interact. Group housed BALB/c males were used as intruders because they are a relatively non-aggressive strain, thereby reducing any intruder-initiated fighting. Behavior videos were hand annotated by an observer blind to experimental conditions (Observer, Noldus).

#### 3-Chamber Sociability Assay

Testing for social preference using the 3-chamber sociability assay proceeded as previously described ^77^. Briefly, experimental mice were placed into a long Plexiglass apparatus (65 × 45 cm) consisting of three equivalently-sized chambers – a center chamber and two side chambers each containing a cup (obtained from the Noldus Sociability Cage) used to separate the conspecifics, either in the top left of the bottom right of the respective chamber, with enough room for the experimental mice to navigate to all sides of the cup. Chambers are divided by white plexiglass walls that include an 11.43cm sized opening to allow mice to freely move between chambers. Mice were allowed to freely explore for a 5-minute baseline period. The baseline phase began once the experimenter removed the doors, allowing the mouse to explore all three empty chambers. After the baseline phase, mice were then ushered back into the middle chamber, with access to other chambers blocked by doors in order to transition to the test phase. The test phase lasted 10 minutes and consisted of an inanimate object (black plastic block) under a cup in one chamber, and a conspecific mouse under a cup in the opposite chamber. The “social” side was counterbalanced for each trial. The apparatus, cups, water bottles, and doors were cleaned with 70% isopropyl alcohol between each trial.

### Behavioral Analysis

Video data from the behavioral assays was captured using (GigE) infrared cameras. Mouse social behavior was scored by an observer blind to the animal’s condition using Noldus annotation software, The Observer. A cadre of social behaviors were manually annotated, including, but not limited to, social investigation, cornering, and fighting. Animal tracking data was collected for the three-chamber sociability assay using Noldus video tracking software, Ethovision. Total time in each chamber, as well as time interacting with the object and mouse cups, was automatically quantified and binned into baseline and testing intervals. The data gathered by video tracking software was confirmed by manual annotation performed by an observer blind to experimental conditions.

### Immunohistochemistry

Mice were transcardially perfused 90 minutes after the mid-point of either the 3Ch-sociabiliity or the RI assay with 4°C phosphate-buffered saline (PBS) followed by 4% paraformaldehyde (PFA) in PBS.

All brains for immunohistochemistry were stored in PFA for 24 hours, then moved to a 15% sucrose solution for 24 hours, and then to a 30% sucrose solution for another 24 hours. Brains were then embedded in Tissue-Tek optimal cutting temperature (OCT) compound and stored at - 80°C until ready to slice. Brains were serial sectioned coronally in 40µm increments using a Leica Cryostat and collected in cold PBS. Slices were rinsed 3 times with 1x PBS for 10 minutes each time at room temperature on an orbital shaker. Sections were then incubated at room temperature on a rocker for 1 hour in blocking solution (10% normal goat serum + 0.3% Triton X-100 in 1x PBS). Sections were then transferred to wells containing 1-2mL of 1:1000 primary antibody (abcam rabbit anti-cfos antibody ab190289) in blocking solution and incubated at 4°C for 48-72 hours. Sections were then rinsed again 3 times in 1 x PBS at room temperature for ten minutes each, then transferred to a 1:250 fluorescent secondary antibody (Invitrogen Goat anti-Guinea Pig Alexa Fluor 594, catalog #A-11076) diluted in blocking buffer and incubated at 4°C overnight. Sections were once again washed 3 times with 1x PBS and then mounted onto positively charged slides, counterstained for DAPI with Vectashield Hardset mounting medium, cover slipped, and left to dry overnight.

### Imaging and Image Analysis

#### cFos Imaging and Quantification

For ventral hippocampal cFos analysis, images were acquired using a Zeiss Axio Scan.Z1 slide scanner within the microscopy core at the University of Utah. Following cFos IHC, mounting, and DAPI staining, slides containing the dorsal and ventral hippocampus were loaded into the slide scanner. A 10x objective was used to take serial images of the dorsal and ventral hippocampus, and these images were stitched together using the Zeiss Imaging software. Images were saved as .czi files and then exported as .tif files using ZEN 3.6 (blue edition) for image analysis. TIF files were then uploaded into QuPath, an open software for bioimage analysis. The region of interest was drawn separately around CA1 in the dorsal and ventral hippocampus. Image gamma was adjusted to remove any background fluorescence, and cFos+ cells were hand-counted by an observer blinded to experimental conditions. The number of positive cells were then divided by the total area (in mm^2^). Ventral hippocampal slices ranging from AP coordinates −2.91 to −3.39 were chosen for vCA1 cell counting.

#### Activity-Dependent Tagging Imaging and Quantification

A Leica DM2000 LED epifluorescent microscope was used to capture tagged cells in the ventral hippocampus. Images were taken using 5x, 10x, and 20x objective lenses. Images were saved as .czi files and then exported as .tif files using ZEN 3.6 (blue edition) for merger and image analysis. All cells tagged during behavior 1 and behavior 2 were hand-counted separately by an observer blinded to experimental conditions. Merged images were used to hand-count overlapping cells (cells active during both behavior 1 and behavior 2). To determine the percentage of overlapping cells, we divided the number of overlapping cells by the number of total cells tagged during each behavior.

#### DREADDs Histological Verification

To confirm viral expression of the DREADDs in the ventral hippocampus, we acquired images using a Leica DM2000 LED epifluorescent microscope with a 5x objective lens. Any animals that had expression outside of the VH were excluded.

### Statistical Analysis

All statistical analyses were performed using GraphPad Prism. Normality tests and respective QQ plots were generated to determine appropriate statistical testing. In most cases the data was normally distributed and parametric testing, such as the t-test (paired/unpaired) and ANOVA (mixed-design/two-way), were performed. However, time spent fighting tends to be non-normally distributed, as a sizable portion of mice do not exhibit this behavior at all. Under these circumstances, non-parametric tests, such as Mann-Whitney, were performed. When appropriate, Sidak’s *post hoc* analyses were conducted. All statistical tests assumed an alpha level of 0.05. Statistical tests are reported in (each figure legend?) with*p<0.05, **p<0.01, and ***p<0.001. Individual data points were presented in the figures, in addition to mean +/- SEM.

## ACKNOWLEDGEMENTS

We thank the Steve Ramirez Lab for providing the viruses necessary for the activity-dependent tagging system (pAAV-cFos-tTA, pAAV-TRE-eYFP). This work was supported by NIMH R00 MH108734 (MZ), NIMH R01 MH132822 (MZ), a Klingenstein-Simons Early Investigator Award (MZ), a Whitehall Fellowship (MZ), a Sloan Fellowship (MZ), a McKnight Scholars Award (MZ), and the University of Utah.

## Notes

### Competing Interest Statement

The authors have declared no competing interest.

