## Supplemental Figures for "Stress induces distinct social behavior states encoded by the ventral hippocampus"

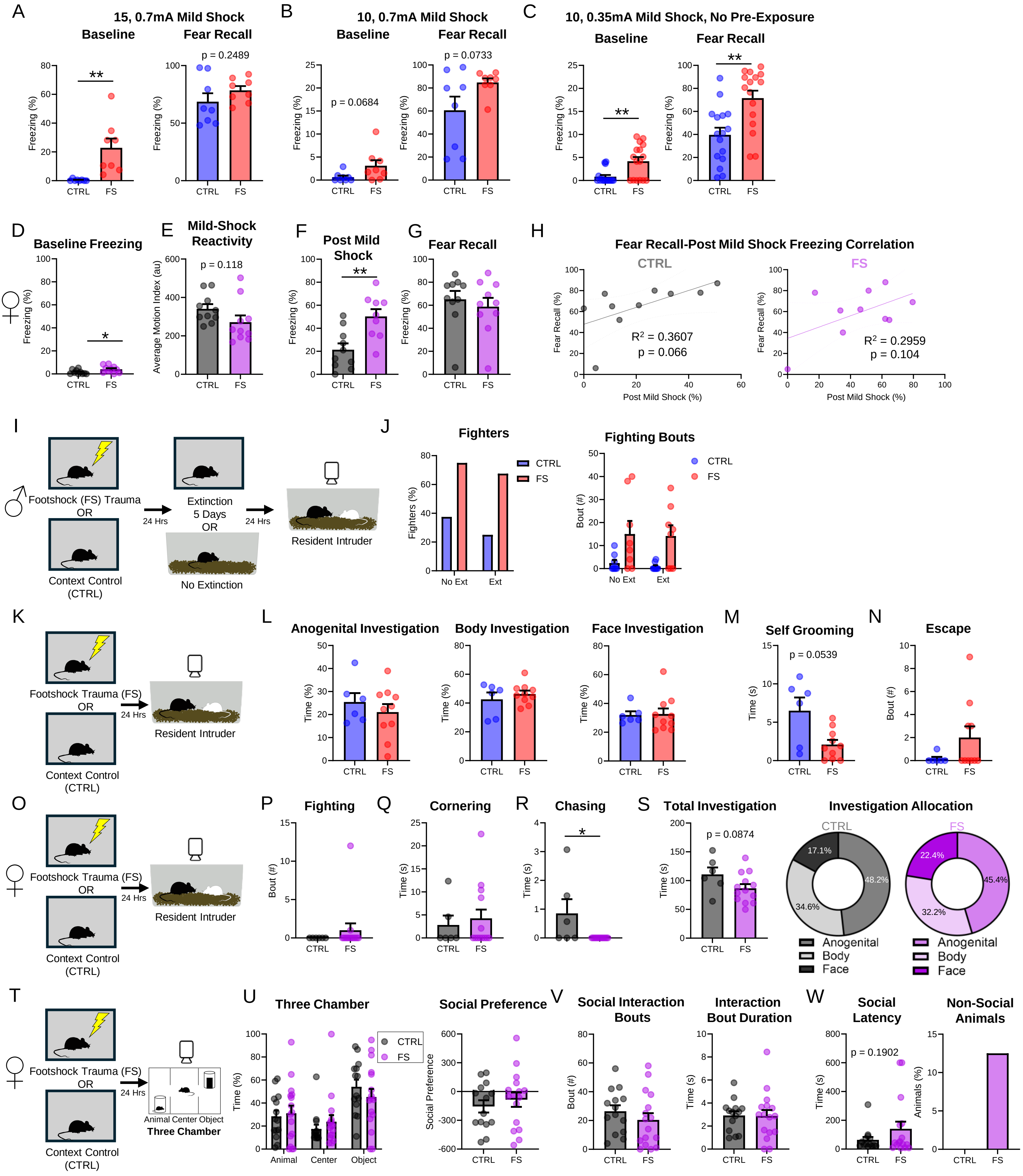
A
15, 0.7mA Mild Shock
Baseline
Fear Recall
p = 0.2489
Freezing (%)
CTRL
FS
B
10, 0.7mA Mild Shock
Baseline
Fear Recall
p = 0.0684
p = 0.0733
C
10, 0.35mA Mild Shock, No Pre-Exposure
Baseline
Fear Recall
D
Baseline Freezing
E
Mild-Shock Reactivity
p = 0.118
Average Motion Index (au)
F
Post Mild Shock
G
Fear Recall
H
Fear Recall-Post Mild Shock Freezing Correlation
CTRL
FS
Fear Recall (%)
Post Mild Shock (%)
R² = 0.3607
p = 0.066
R² = 0.2959
p = 0.104
I
Footshock (FS) Trauma
OR
Context Control
(CTRL)
24 Hrs
Extinction
5 Days
OR
No Extinction
24 Hrs
Resident Intruder
J
Fighters
Fighters (%)
No Ext
Ext
Fighting Bouts
Bout (#)
K
Footshock Trauma (FS)
OR
Context Control
(CTRL)
24 Hrs
Resident Intruder
L
Anogenital Investigation
Body Investigation
Face Investigation
Time (%)
M
Self Grooming
p = 0.0539
Time (s)
N
Escape
Bout (#)
O
Footshock Trauma (FS)
OR
Context Control
(CTRL)
24 Hrs
Resident Intruder
P
Fighting
Q
Cornering
R
Chasing
S
Total Investigation
p = 0.0874
Investigation Allocation
CTRL
FS
17.1%
48.2%
34.6%
22.4%
45.4%
32.2%
Anogenital
Body
Face
T
Footshock Trauma (FS)
OR
Context Control
(CTRL)
24 Hrs
Animal Center Object
Three Chamber
U
Three Chamber
Animal
Center
Object
Social Preference
V
Social Interaction Bouts
Interaction Bout Duration
W
Social Latency
p = 0.1902
Non-Social Animals
Animals (%)

**Supplemental to Figure 1.**
(A) – (C) Differences in baseline fear and fear recall across different shock parameters.
(D) Female baseline freezing.
(E) Female mild-shock reactivity.
(F) Female post mild footshock.
(G) Female fear recall.
(H) Female correlation between post-shock and fear recall test freezing in CTRL (left) and FS (right) mice.
(I) Experimental design to test for fighting behavior following fear extinction.
(J) Percent male fighters between CTRL and FS with and without fear extinction training (left) and number of fighting bouts (right).
(K) Experimental design to examine behaviors during resident intruder (males).
(L) Male individual investigation types.
(M) Male self-grooming behavior.
(N) Male escape behavior.
(O) Experimental design to examine behaviors during resident intruder (females).
(P) Female fighting bouts.
(Q) Female cornering time.
(R) Female chasing time.
(S) Female total investigation time (left) and investigation allocation (right)
(T) Experimental design to examine behaviors during three-chamber sociability assay (females).
(U) Female percentage of total time spent in each chamber (left) and social preference score (right).
(V) Female social interaction bouts (left) and interaction bout duration (right)
(W) Female social latency (left) and percentage of animals that never approached the social chamber (right).
Data are presented as mean ± SEM. Significant differences are reported as ****$p < 0.0001$, ***$p < 0.001$, **$p < 0.01$, and *$p < 0.05$.

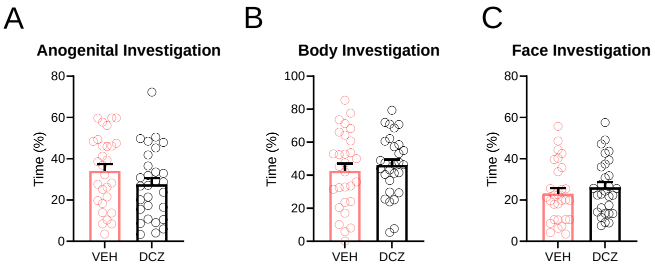
A
Anogenital Investigation
Time (%)
VEH
DCZ
B
Body Investigation
Time (%)
VEH
DCZ
C
Face Investigation
Time (%)
VEH
DCZ

**Supplemental to Figure 3. Investigation allocation comparing male mice with ventral hippocampus silenced vs those with ventral hippocampus active.**
(A) Male anogenital investigation allocation.
(B) Male body investigation allocation.
(C) Male face investigation allocation.
Data are presented as mean ± SEM. Significant differences are reported as ****$p < 0.0001$, ***$p < 0.001$, **$p < 0.01$, and *$p < 0.05$.
